# Characterization of a new *Leishmania major* isolate for use in a controlled human infection model

**DOI:** 10.1101/2020.08.12.245936

**Authors:** Helen Ashwin, Jovana Sadlova, Barbora Vojtkova, Tomas Becvar, Patrick Lypaczewski, Eli Schwartz, Elizabeth Greensted, Katrien Van Bocxlaer, Marion Pasin, Kai S. Lipinski, Vivak Parkash, Greg Matlashewski, Alison M. Layton, Charles J. Lacey, Charles L. Jaffe, Petr Volf, Paul M. Kaye

**Author notes:** These authors contributed equally to this study (Helen Ashwin, Jovana Sadlova).

## Abstract

Leishmaniasis is widely regarded as a vaccine-preventable disease, but the costs required to reach pivotal Phase 3 studies and uncertainty about which candidate vaccines should be progressed into human studies significantly limits progress in vaccine development for this neglected tropical disease. Controlled human infection models (CHIM) provide a pathway for accelerating vaccine development and to more fully understand disease pathogenesis and correlates of protection. Here, we describe the isolation, characterization and GMP manufacture of a new clinical isolate of *Leishmania major*. Two fresh isolates of *L. major* from Israel were initially compared by genome sequencing, *in vivo* infectivity and drug sensitivity in mice, and development and transmission competence in sand flies, allowing one (*L. major*_MRC-02) to be selected for GMP production. This study addresses a major roadblock in the development of vaccines for leishmaniasis, providing a key resource for CHIM studies of sand fly transmitted cutaneous leishmaniasis.

## Introduction

The leishmaniases represent a group of diseases caused by infection with various species of the parasitic protozoan *Leishmania*. One billion people are at risk of infection across 98 countries worldwide, with over 1.5 million new cases and 20,000 – 40,000 deaths reported each year ^1,2^. The leishmaniases are vector-borne diseases, each parasite species having co-evolved for transmission by one or more species of phlebotomine sand fly ^3,4^. Disease may be evident as self-healing lesions restricted to the site of skin transmission (cutaneous leishmaniasis; CL), lesions which spread from an initial skin lesion to involve the mucosae (mucosal leishmaniasis; ML) or which spread uncontrolled across the body (disseminated or diffuse cutaneous leishmaniasis; DCL), or as a potentially fatal systemic disease involving major organs such as the spleen, liver and bone marrow (kala azar or visceral leishmaniasis; VL) ^5^. In addition, patients recovering from VL following chemotherapy often develop a chronic skin condition (post kala-azar dermal leishmaniasis; PKDL) that can sustain community transmission of VL ^6,7^. Collectively, the tegumentary forms of leishmaniasis account for approximately two-thirds of the global disease burden, whilst VL accounts for most reported deaths ^1^. In addition to the impact of primary disease, recent studies have also emphasized the importance of considering the long term sequelae of leishmaniasis, notably those associated with stigmatization, when evaluating global burden of these diseases ^8–10^.

The leishmaniases are widely regarded as a vaccine-preventable diseases, based on disease natural history, epidemiological data and studies in experimental models of leishmaniasis (reviewed in ^11–14^. Although vaccines for canine visceral leishmaniasis have reached the market, to date no human vaccines have achieved licensure ^15^. Often cited barriers to vaccine development include limited investment, with only $3.7M of new R&D funding globally in 2018 ^16^, an excess of candidate antigens and delivery systems ^17^, the questionable predictive capacity of pre-clinical animal models ^18,19^, lack of good correlates of protection and a defined target product profile ^20^, the costs and challenge of large-scale efficacy studies in disease endemic countries and a fragmented pipeline for translational research ^21^. As has been found with other diseases ^22–29^, the incorporation of a controlled human infection model (CHIM) into the vaccine R&D pipeline can overcome many of these issues.

Artificial human infection (“leishmanization”) with *Leishmania* had been practiced for centuries by people living in the Middle East and former Soviet states, where CL is highly endemic. Building on the knowledge that cure from CL engendered protection against reinfection, scrapings from active lesions were used to cause disease at a site of choice (e.g. the buttock), so avoiding the stigmatization associated with CL scars (reviewed in ^30^. More defined experimental studies were conducted sporadically through the 20^th^ century, culminating in a WHO-sponsored evaluation of the potential for human challenge as a tool to evaluate vaccines for leishmaniasis, conducted in Iran in 2005 ^31^. This study employed a *L. major* isolate that had been produced at GMP and used for previous leishmanization studies. Results from this study demonstrated a take rate of 86% in previously non-exposed volunteers. Lesions were less than 3cm diameter and ulcerated in 74% of cases. All lesions self-healed without treatment between 75 and 285 days after inoculation. In a limited re-challenge study using the same isolate, 0/11 volunteers receiving leishmanization developed a lesion, compared to 5/5 in non-leishmanized controls ^31^. Poor viability of the challenge agent and limited funding opportunities curtailed this program before it could be developed further and to date, no defined vaccines have been tested using this approach.

With the advent of new candidate vaccines in or approaching the clinic, there is renewed imperative to develop a CHIM for leishmaniasis. An adenoviral-vectored vaccine (ChAd63-KH) was found to be safe and immunogenic in healthy volunteers ^32^ and in PKDL patients (Younnis et. al. submitted) and is currently in Phase IIb as a therapeutic in Sudanese PKDL patients. A live genetically-attenuated *L. donovani* centrin^−/−^ parasite has shown efficacy in pre-clinical models ^33–35^ and a *L. major* centrin^−/− 36^ is soon to enter GMP production. An adjuvanted recombinant polyprotein vaccine (LEISH-F3 / GLA-SE) has been progressed to Phase I ^37^ and a newer derivative (LEISH-F3+ / GLA-SE) evaluated in pre-clinical models ^19^. RNA-based vaccines are also in development ^38^. In addition, new knowledge regarding the integral nature of sand fly transmission to *Leishmania* infectivity has emerged in recent years ^39–41^, underpinning the observation that vaccines inducing protection in mice when infected via needle inoculation fail to protect against sand fly transmitted infection ^42^; hence the need to incorporate vector transmission as part of a CHIM.

The pathway for development of a CHIM for sand fly-transmitted leishmaniasis requires three enabling activities: the identification of an appropriate challenge agent, optimization of sand fly transmission studies in humans, and patient and public involvement (PPI). Here, we describe completion of the first of these steps, namely the isolation, characterization and GMP production of a new *L. major* challenge agent.

## Results

### New clinical isolates of Leishmania major

*Leishmania major* is endemic in Israel and cases are often associated with travelers visiting areas of high transmission ^43,44^. Two individuals from non-endemic areas of central Israel that had self-referred to Sheba Hospital in early 2019 after developing lesions subsequent to visiting the endemic region of Negev (**Figure 1A**) served as parasite donors. Donor MRC-01 was a forty-one-year-old female who developed a lesion near her left lip approximately three months after spending one night outdoors. She had self-administered topical antibiotics without effect and presented at clinic approximately 4 months later with a single erythematous 1.5cm diameter lesion. Diagnosis for *L. major* was confirmed by PCR and she was treated with intra-lesional sodium stibogluconate (SSG) on two occasions approximately 4 weeks apart. Her lesion fully resolved with minimal scarring by 3 months post treatment onset (**Figure 1B**). Donor MRC-02 was a twenty-two-year-old male who developed two papules on his shin approximately two months after hiking in the Negev. He attended clinic three months later with two ~1.5cm diameter ulcerated lesions on the shin and a very small non-ulcerated lesion on his neck. He was diagnosed positive for *L. major* by PCR but refused treatment. His lesions fully resolved approximately three to four months later with scarring (**Figure 1C**). Both donors were negative for HIV, HTLV-1, HBV and HCV, and at 18-month follow up, neither reported any reactivation of their lesion(s) or other unexpected clinical events related to their leishmaniasis.

**Figure 1.**
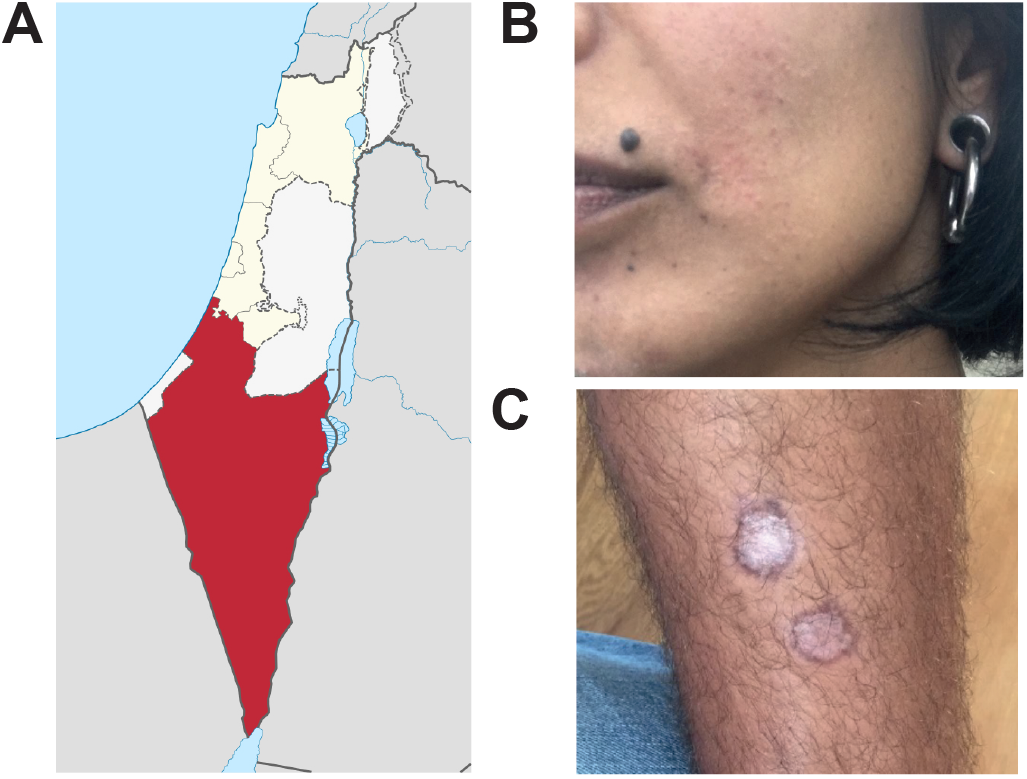
Clinical characteristic of patient lesions. **A.** Negev region of Israel (red). **B.** Donor MRC-01 lesion photographed ~3 months after treatment onset, showing full resolution of the lesion near the corner of the mouth, with minimal scarring. **C.** Donor MRC-02 lesions photographed ~9 months after travel to the endemic region, showing healing without treatment.

### Whole genome sequencing of parasite isolates from patients MRC-01 and MRC-02

Parasites were isolated from slit skin smears and diagnosis of *L. major* infection confirmed by PCR and RFLP analysis. The isolates were designated as *L. major*_MRC-01 and *L. major*_MRC-02, respectively. Parasites were minimally cultured in GMP grade media to retain infectivity and multiple vials frozen at P1 as a seed bank and screened negative for mycoplasma. The seed stock was redistributed under dry ice using a commercial shipping agent.

To confirm genetic identity and establish baseline sequence data, whole genome sequencing was performed using Illumina NextSeq deep sequencing. Sequence data for *L. major*_MRC-01 and *L. major*_MRC-02 has been deposited at GenBank. A phylogeny tree was developed using all available whole genome *L. major* sequences from around the world with *L. major* Friedlin strain from Israel used as reference (**Figure 2A**). This analysis shows a geographical clustering of genome sequences and confirms that *L. major*_MRC-01 and *L. major*_MRC-02 are closely related to other *L. major* isolates derived from Israel while being distinct from each other. Next, sequence alignment was performed to identify the location of single nucleotide polymorphisms relative to the reference Friedlin strain, using DNA from early culture passage *L. major*_MRC-01 and *L. major*_MRC-02 (grey lines, **Figure 2B**). As shown in the inner rings, both MRC-01 and MRC-02 isolates had a number of SNPs compared to the reference Friedlin strain in all 36 chromosomes (**Figure 2B**, rings MRC01Pre and MRC02Pre). Similarly, the SNP fingerprint of MRC-01 compared to the MRC-02 isolate also reveal these are genetically distinct isolates consistent with the phylogeny tree.

**Figure 2:**
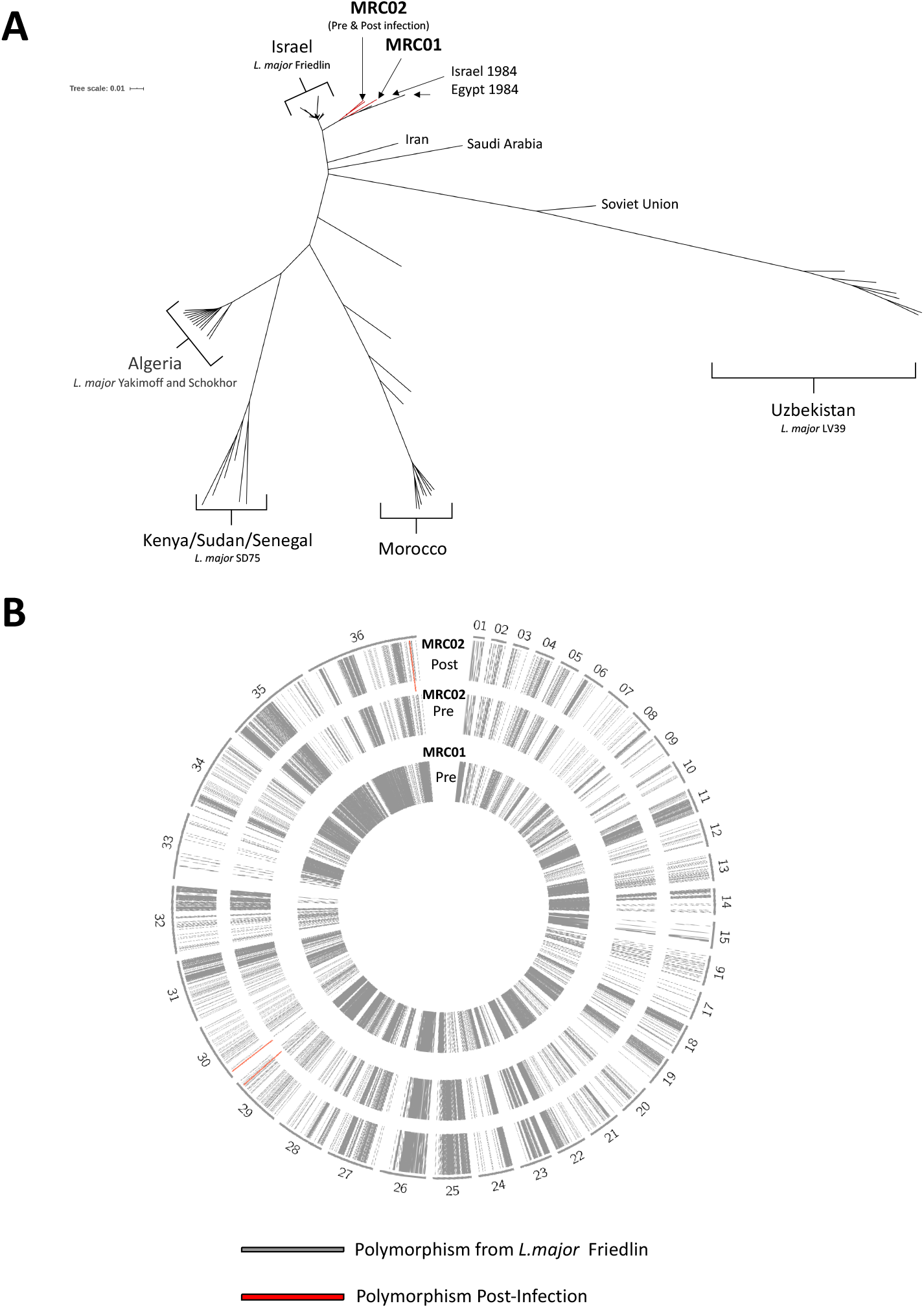
Characterisation of *L. major*_MRC-01 and *L. major*_MRC-02 by whole genome sequencing. **A.** Phylogeny tree developed using all available whole genome sequences for *L. major* from different parts of the world and the location of the MRC-01 and MRC-02 isolates. The phylogeny tree was constructed using the *L. major* Friedlin strain as the reference strain. **B.** Alignment map for *L. major* chromosomes 1 – 36. Grey bars represent location of single nucleotide polymorphism (SNPs) and indels where there are differences between the *L. major* MRC-01 and *L. major* MRC-02 and *L. major* MRC-02 (Post: following passage in BALB/c mice) and the reference strain *L. major* Friedlin. Red bars indicated chromosomal location of SNPs/indels differences between *L. major* MRC-02 (Pre-passage) and the *L. major* MRC-02 (Post-passage).

We also compared whole genome sequences from *L. major*_MRC-02 before and after a single passage in BALB/c mice (see below). Compared to early culture parasites (pre-infection), the SNP fingerprint of the BALB/c passaged parasites (post-infection) was nearly identical when comparing the 2 outer rings (**Figure 2B,** rings MRC02-Post, MRC02-Pre). Only three polymorphisms were identified in parasites recovered from *in vivo* passage, and all were in homopolymer stretches in non-coding regions (**Table S1A and S1B**). In addition, we found no significant copy number variation (CNV) differences at the gene or chromosome level between the genomes from *L. major*_MRC-02 (Pre, culture) and *L. major*_MRC-02 (Post, *in vivo* passage) parasites (**Figure S2**). These observations confirm the *L. major-_*MRC-01 and *L. major* MRC-02 isolates are closely related to strains previously isolated from Israel, are genetically distinct from each other and that there was no selection for genomic mutations or CNVs following MRC-02 infection in BALB/c mice.

*Leishmania* RNA viruses (LRV) have been demonstrated in various *Leishmania* species: LRV1 in *L.* (*Viannia*) *braziliensis* and *L. (V.) guyanensis*, and LRV2 in *L. aethiopica, L. infantum, L. major* and *L. tropica*. RNA was isolated from early passage *L. major*_MRC01 and *L. major*_MRC02, and tested for LRV2 by RT-PCR ^45^. *L. aethiopica* LRC-L494, previously shown to contain LRV2, was used as a positive control. Both *L. major* isolates were negative for LRV2 (**Figure S3**).

### In vitro and in vivo characterization and drug sensitivity of L. major_MRC01 and L. major_MRC02

Prior to *in vivo* infectivity studies, each line was evaluated for growth under standard *in vitro* conditions. Both lines showed similar *in vitro* growth curves, with characteristic progression through logarithmic and stationary phases of growth (**Figure 3A**). Metacyclic promastigotes (**Figure 3B**) were isolated by negative selection using PNA ^46^ and used for infectivity studies in mice. To confirm in vivo infectivity and assess sensitivity to paromomycin (PM), a standard drug used for the treatment of CL ^47^, we used highly susceptible BALB/c mice infected subcutaneously in the rump with ~10^6^ purified metacyclic promastigotes (**Figures 3 and 4**). Mice were randomized to receive PM (50mg/kg i.p. daily for 10 days), with treatment starting when individual lesion size was 3-4mm in diameter. Both isolates induced lesions in BALB/c mice (**Figures 3C and E**). In the absence of treatment, all infected mice progressed to the pre-determined endpoint (8mm average diameter, <10mm in any direction) or showed progressing disease at the experimental endpoint of 70 days post infection. However, *L. major*_MRC-02 lesions developed more rapidly and in a more consistent manner. For example, median time to develop a lesion > 2mm was 37.5 vs. 21.0 days, for *L. major*_MRC-01and *L. major*_MRC-02 respectively (ratio 1.786, 95% CI of ratio 0.96 to 3.32; p<0.0001, **Figures 3D-G** and **Figure S4**). All mice responded well to PM treatment, with a reduction in lesion size that often reached the limits of detection within the 10-day treatment window (**Figures 3E** and **G**, and **Figures 4A-D**). Real time PCR quantification of parasite kDNA in the lesion indicated that PM treatment reduced parasite load for both isolates by >99% (**Figure 4E**). Thus, whilst both isolates are capable of causing lesions in BALB/c mice and can be cured using PM, *L. major*_MRC-02 demonstrates more rapid lesion development, with greater reproducibility.

**Figure 3.**
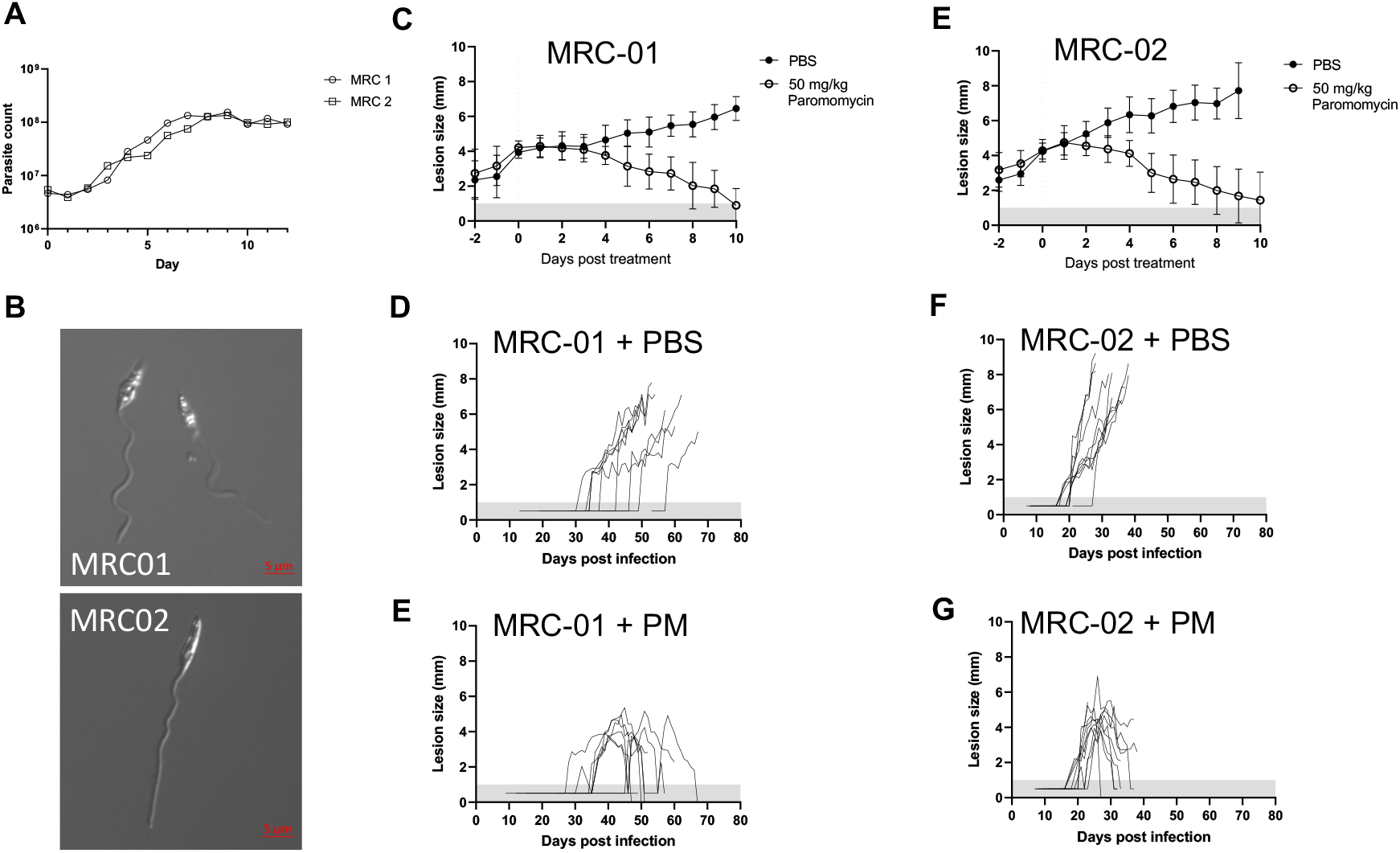
Growth characteristics and drug sensitivity of *L. major*_MRC-01 and *L. major*_MRC-02. **A.** Growth curves for *L. major*_MRC-01 and *L. major*_MRC-02 in vitro. **B.** Photomicrographs of purified PNA-negative metacyclics of *L. major*_MRC-01 (top) and *L. major*_MRC-02 (bottom). **C-G**. In vivo lesion development in BALB/c mice following subcutaneous infection with 10^6^ metacyclics of *L. major*_MRC-01 or *L. major*_MRC-02 in the presence or absence of 50mg/kg paromomycin (PM) i.p. daily for 10 days. Treatment was initiated when lesion diameter reached 3-4mm and mice were killed if lesion size exceeded 9-10mm in any one direction. Data are presented in aggregated form (C, E) normalised to the day of treatment initiation (shown as dotted vertical line) and as a timeline for individual mice receiving vehicle alone (D, F) or PM treatment (E, G). Data are derived from two independent experiments with 9-10 mice per group per strain. Data points within horizontal shaded area represent lesion was palpable but not measurable at <1mm diameter.

**Figure 4.**
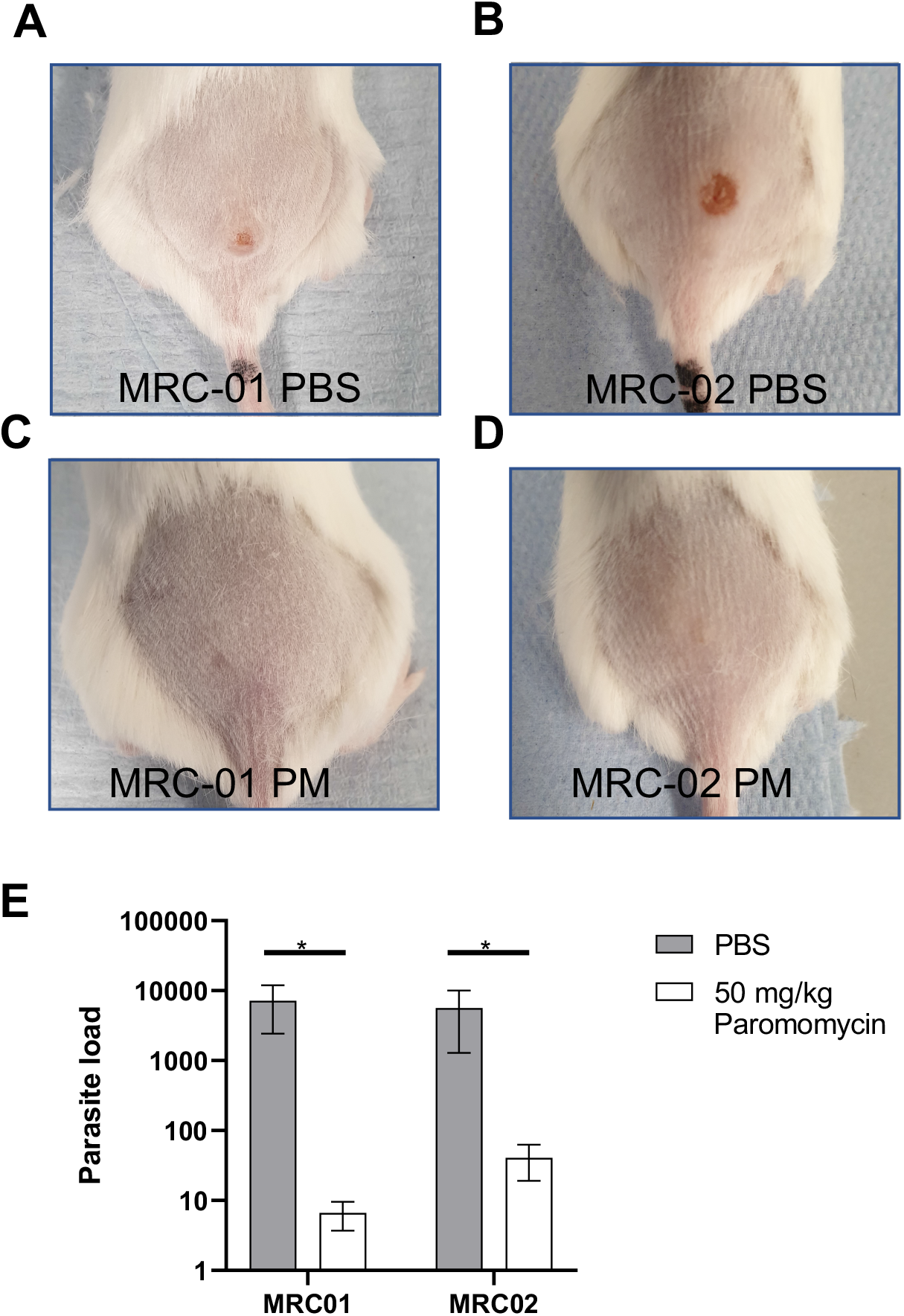
Response of BALB/c mice infected with *L. major*_MRC-01 and *L. major*_MRC-02 to paromomycin treatment. **A-D.** Representative photographs of BALB/c mice infected with *L. major*_MRC-01 (A, C) and *L. major*_MRC-02 (B, D) in absence of treatment (A, C) and at the end of 10 days PM treatment (B, D). **E.** Parasite loads in lesions from PM treated and untreated mice, as determined by qPCR. Data are derived from n=9-10 mice per group per strain and from two independent experiments

### Parasite development in the sand fly vector

To determine whether *L. major*_MRC01 and *L. major*_MRC02 were fully competent for sand fly transmission, we first conducted artificial membrane feeding experiments using two vector species, *Phlebotomus papatasi* and *P. duboscqi*. Experimental infections indicated that both isolates developed well in the two sand fly species (**Figure 5**), producing high infection rates (100% of sand fly females infected by day 3 post blood meal (PBM), >75 % at days 6 and 15 PBM). In *P. duboscqi*, development was more vigorous, with parasite escape from the peritrophic matrix and colonization of the thoracic midgut, cardia and in some cases the stomodeal valve by day 3 PBM. In contrast, in *P. papatasi* the first colonization of the stomodeal valve was not observed until day 6 PBM (**Figure 5B**). Nevertheless, by day 15 PBM, both sand fly species had supported full development of parasites, with heavy parasite loads in the thoracic midgut and colonization of the stomodeal valve in all the female sand flies infected with both *L. major* isolates.

**Figure 5.**
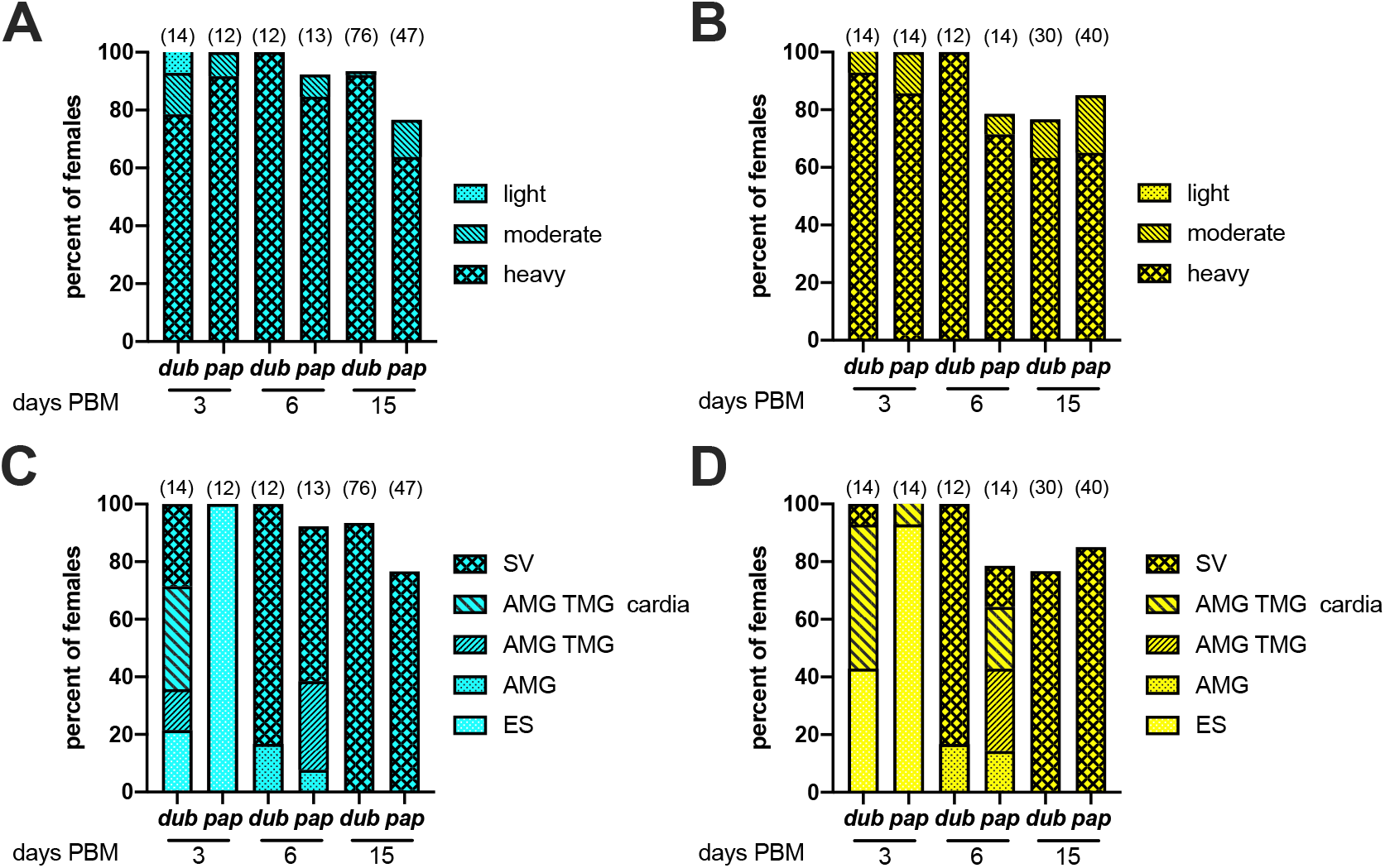
Qualitative analysis of *L. major*_MRC-01 and *L. major*_MRC-02 development in sand flies<. At the days indicated post blood meal (PBM), engorged *P. duboscqi* (*dub*) and *P. papatasi* (*pap*) were dissected and parasite development was assessed. **A and B.** Percentage of flies infected with *L. major*_MRC-01 (A) and *L. major*_MRC-02 (B) based on assessment of intensity of infection. **C and D.** Development of *L. major*_MRC-01 (C) and *L. major*_MRC-02 (D) as assessed by location within the gut; endoperitrophic space (ES), anterior midgut (AMG), thoracic midgut (TMG) cardia and stomodeal valve (SV). Data are pooled from three independent experiments and are shown as frequency of total number of sand flies dissected. Number of sand flies dissected is shown above each bar.

To more precisely quantify parasite load at each morphological stage, exact numbers of procyclic and metacyclic forms in infected sand fly females were counted using a Burker chamber. The differences between *Leishmania* isolates were not significant, indicating that both vectors were capable of supporting development. There was a trend for greater numbers of metacyclic parasites in sand flies infected with *L. major*_MRC-02 at day 3 PBM, but this was not apparent by day 15 PBM, with metacyclic numbers ranging from 200 – 258,000 per sand fly (**Figure 6** and **Table S1**). Given recent data suggesting that additional blood meals may serve to enhance the development of metacyclics ^41^, we conducted a pilot experiment in which we provided sand flies either one additional blood meal on an uninfected BALB/c mouse at day 6 or two additional blood meals at day 6 and day 12 PBM. Under the conditions used, we found no significant differences in metacyclic numbers in *P. duboscqi* infected with either *L. major* isolate using these different feeding conditions (**Table S1**). Although both vector species could therefore be suitable for use in a CHIM, the additional robustness of *P. duboscqi*, and a trend towards more permissive parasite development (^48^ and this manuscript) favor use of this species.

**Figure 6.**
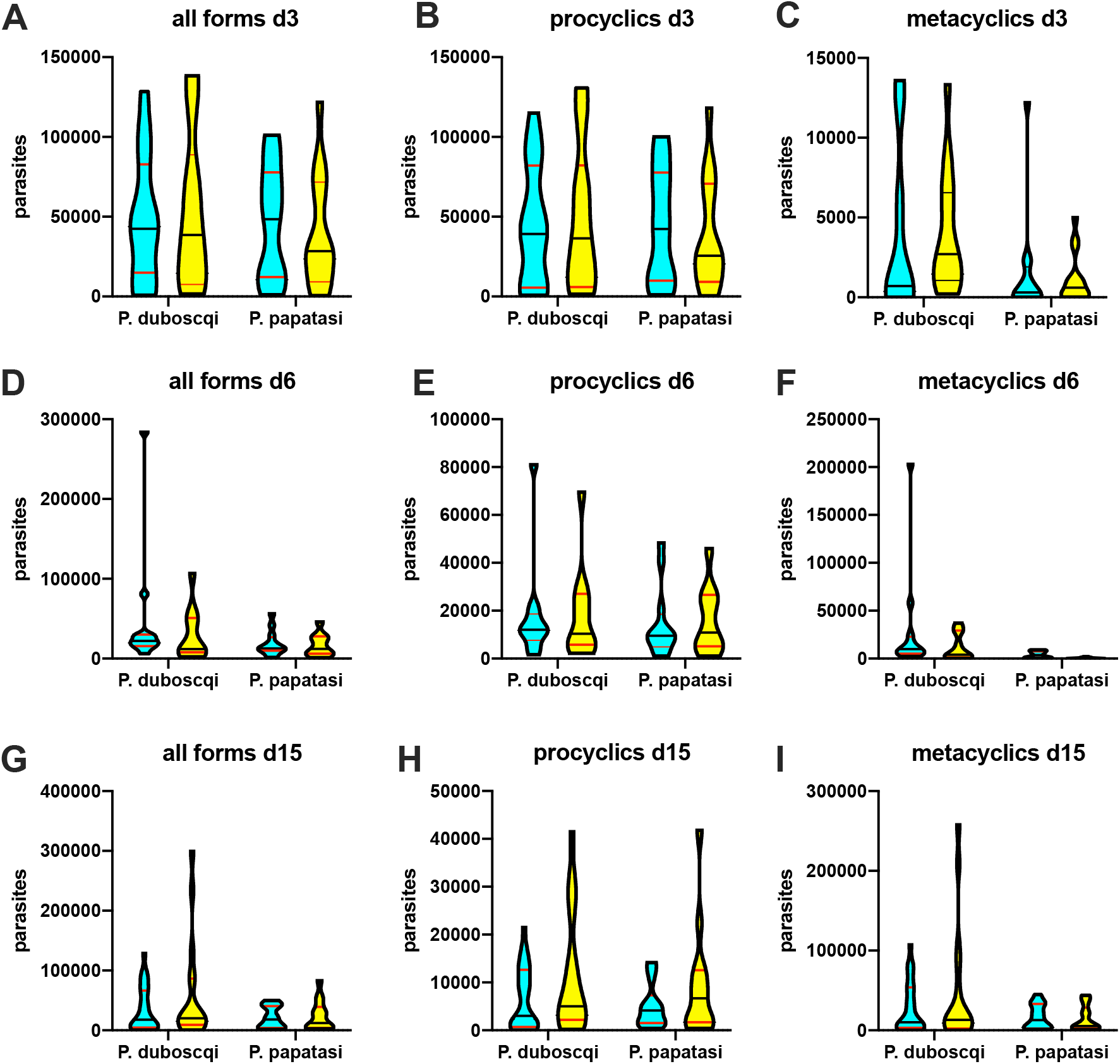
Quantitative analysis of *L. major*_MRC-01 and *L. major*_MRC-02 development in sand flies. At the days indicated PBM, the number of *L. major*_MRC-01 (blue) and *L. major*_MRC-02 (yellow) parasites in engorged sand flies was quantified. **A-C.** Parasite loads in sand flies at day 3 PBM. **D-E**. Parasite loads in sand flies at day 6 PBM. **F-G.** Parasite loads in sand flies at day 15 PBM. Data are shown for all parasites (all forms, A, D, E) and separately for procyclics (B,E,H) and metacyclics (C,F,I). Data are presented as violin plots truncated at the max/min values, with median (black line) and quartiles (red line) indicated and reflect counts obtained from 9-25 individual sand flies of each species dissected per time point for each infection. Raw data can be found in **Table S1**.

To evaluate whether expansion under GMP conditions might affect parasite development, we repeated these experiments using parasites expanded as a research bank (RB) under conditions identical to that for proposed GMP manufacture. Given the data above, we limited these experiments to *P. duboscqi* given a single infectious blood meal by membrane feeding. As before, both *L. major* isolates produced 100% infection rates in sand fly females on day 3 PBM, with >80 % late stage infections on day 6 and day 15 PBM. At day 3 PBM, 14% and 43% respectively of females infected with *L. major*_MRC-01 and *L. major*_MRC-02 had parasites located at the stomodeal valve. By day 15 PBM, thoracic midguts were filled with high numbers of parasites and the stomodeal valve was colonized in all the females infected with both isolates, though heavier infections developed in sand flies infected with *L. major*_MRC-02 (**Figure S5)**. Thus, the limited expansion required to generate a GMP parasite bank does not negatively impact on parasite development in sand flies.

### Transmission of L. major_MRC01 and L. major_MRC02 to mice by sand fly bite

Ten *P. duboscqi* females infected by *L. major*_MRC-01 or *L. major*_MRC-02 were allowed to feed on anaesthetized BALB/c mice on day 15 post BM. Immediately post feeding, six ear samples per each strain were taken for determination of transmitted parasite number using qPCR. Positivity rates were 5/6 and 6/6 for *L. major*_MRC-01 and *L. major*_MRC-02, respectively, and numbers of parasites per ear varied from 0 to 7240 and from 92 to 5670 respectively for the two isolates (**Figure 7A** and **Table S2**). The average numbers of transmitted parasites did not differ significantly between the two groups.

**Figure 7.**
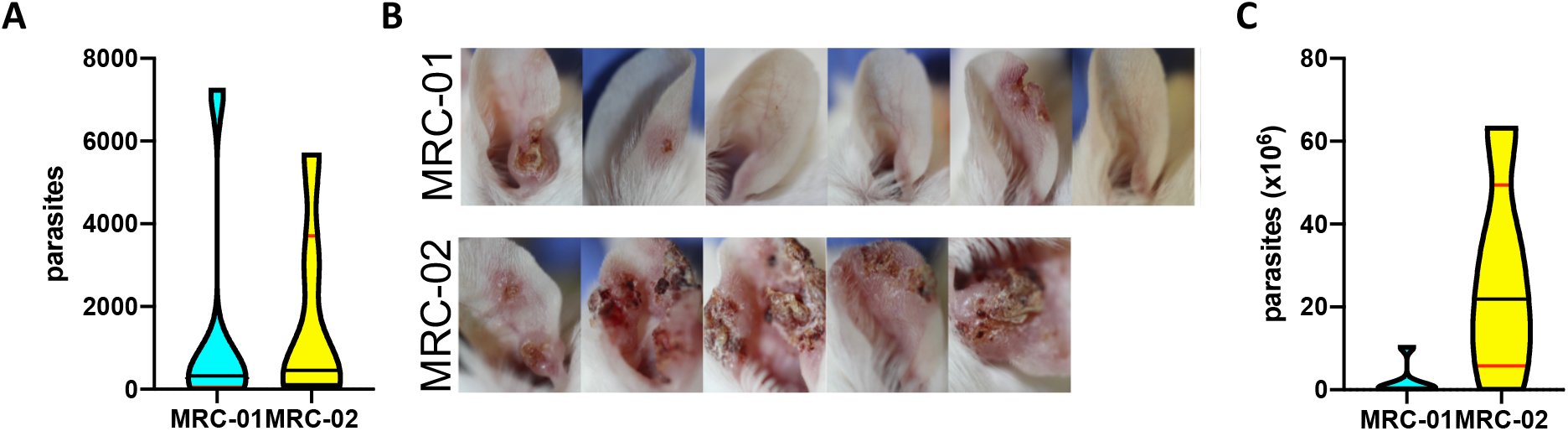
Transmission of *L. major*_MRC-01 and *L. major*_MRC-02 to BALB/c mice by sand fly bite. **A.** Parasites loads per ear determined by qPCR immediately post bite. Data is shown as violin plot for n=6 ears per parasite strain. Additional data found in Table S3. **B.** Photographs of ear lesions in individual mice 6 weeks post exposure to sand flies infected with *L. major*_MRC-01 and *L. major*_MRC-02. For time course photographs, see **Figure S6**. **C**. Parasite loads determined at 6 week post exposure to infected bites. Data is shown as violin plot for n=5 (MRC-02) or 6 (MRC-01) ears per parasite strain.

An independent group of mice was followed to monitor lesion development. Three weeks p.i., only ear swelling was observed in mice bitten by sand flies infected with *L. major*_MRC-01, whereas lesions had developed in 50% (3/6) of mice exposed to sand flies infected with *L. major*_MRC-02. At the end of the experiment on week 6 p.i., all five mice bitten by *L. major*_MRC-02 infected flies showed presence of skin lesions while lesions appeared only in a half of mice (3/6) bitten by *L. major*_MRC-01 infected flies (**Figure 7B, Table S2** and **Figure S6**. Determination of parasite load by qPCR at 6 weeks p.i. indicated a trend towards higher numbers of parasites in mice infected with *L. major*_MRC-02 (P = 0.08, **Figure 7C**). Of note, two mice exposed to sand flies infected with *L. major*_MRC-01 hosted significant numbers of parasites in the ear (2.82×10^5^ and 6.44×10^5^) despite the absence of lesions (**Table S2**). Hence, *L. major*_MRC-02 produces rapid and reproducible lesions in mice after sand fly transmission.

### *GMP Production of L. major*_MRC-02

A GMP clinical lot of *L. major*_MRC-02 was produced under contract directly from P1 passage stocks using static T-flask cultures. The clinical lot comprises ~600 vials, each vial containing 2×10^7^ mid log *L. major*_MRC-02 in culture media. Vials are stored at −145 ± 10°C and we expect shelf life to exceed 5 years. An initial 2 years stability study will be performed at Vibalogics manufacturing site. Release testing of the batch was discussed and agreed with the UK Medicines and Healthcare products Regulatory Agency (MHRA) and comprises identity (PCR), resuscitation (indicating growth), sterility, endotoxin and pH. We estimate conservatively that in a sand fly CHIM, after retention for stability studies, this clinical lot will be sufficient for challenge of at least 1200 volunteers. Seed stocks are available for further GMP runs as required.

## Discussion

Controlled human challenge is increasingly viewed as being on the critical path for vaccine development, allowing an early demonstration of efficacy and in combination with appropriately designed Phase I trials, rapid selection / de-selection of candidate vaccines ^49^. We have therefore sought to develop a new CHIM, based on best practices derived from other models. Three questions influenced our approach to developing a challenge agent, namely which: i) parasite species, ii) challenge route and iii) manufacturing standard?

Addressing the first question, *L. major*, the causative agent of Old World CL lends itself to development as a human challenge agent on a number of counts. First, unlike other species causing CL e.g. *L. tropica*, *L. mexicana* or *L.* (*Viannia*) *braziliensis* and *L*. (*V*.) *guyanensis*, where systemic or metastatic spread is commonly documented, lesion development following *L. major* infection is usually localised to the site of sand fly transmission and is most commonly self-healing ^5^. Numerous barriers to developing a CHIM model of CL using existing *L. major* isolates were identified, including limited information on the provenance of parasites held in depositories or in use in research laboratories. Although CHIM studies per se are not under formal regulatory control in the UK, informal advice from the MHRA emphasised the importance of understanding the clinical history of the challenge agent donor, donor status with regard other human infectious agents (e.g. HIV) and the need to ensure absence of contact with bovine sera potentially contaminated with agents known to cause transmissible spongiform encephalopathies. In the case of *Leishmania*, passage history also represents an additional, but often poorly defined, variable that governs infectivity ^50,51^. Hence, two fresh isolates were obtained from donors with documented clinical histories and for which we could ensure complete traceability of culture history.

We conducted whole genome sequencing to establish baseline characteristics and to assess genetic changes occurring after passage in animals. The two parasite isolates we examined were genetically distinct, but no features were identified that directly pertain to their potential value as a CHIM agent, given that relatively little is known about the genetic nature of virulence in *Leishmania* parasites. Whilst virulence factors / pathways have been identified in various species, including gp63, lipophosphoglycan, exosome production, proteases and many others ^52^, how these vary across species or isolates and associate with different clinical presentations is poorly defined ^53^. Symbiotic leishmaniaviruses have been associated with enhanced host type I interferon responses and contribute to the metastatic potential in *L. Viannia* species ^54^. Although *Leishmaniavirus* has been detected at lower frequency in Old World *L. major* strains ^55,56^, there is no conclusive data to suggest an involvement in pathogenesis and treatment failure. In Iranian cases, *L. major* infection was not influenced by the presence or absence of LRV2 ^57^. In any event, both *L. major*_MRC-01 and *L. major*_MRC-02 were demonstrated to be negative for LRV2.

The classical BALB/c mouse model was used to evaluate *in vivo* infectivity and drug sensitivity to paromomycin, an often used drug for the treatment of CL ^47,58^. Parasite development in two species of sand fly, both natural vectors of *L. major* confirmed full life cycle completion, including transmission to mice. The latter experiments also pertain directly the second question, route of challenge agent delivery. Vector transmission has been performed for other CHIMs ^29,59^, and though this approach introduces potential confounding factors such as variability in infectious dose and “take rate” compared to needle injection, this needs to be weighed against the added value of such a model. For leishmaniasis, the body of evidence indicating a synergistic role of sand fly salivary components and parasite secretory products in promoting infectivity ^39,40,60,61^ coupled with the value of sand fly challenge in identifying vaccine candidates ^18,19,42,62^ together make a compelling case to proceed with a natural challenge model.

Finally, we considered whether manufacture should be to GMP or GMP-like, as proposed by others. As it is not possible to generate sand flies that are “GMP”, it might have been argued that GMP-like would be sufficient for our purposes. However, given the limited additional financial costs of producing a clinical lot to GMP and that this leaves open the possibility for direct needle challenge or for any future change in the regulatory framework, we chose to adopt full GMP production of the clinical lot.

We selected *L major_* MRC-02 as the challenge agent for GMP production based on an assessment of risk vs reproducibility for participants enrolled in future CHIM studies. Although *L major_* MRC-02 appeared to develop more aggressive lesions in mice, reproducibility of take and more rapid lesion development significantly decreases the number of participants required and duration of a clinical trial. For example, in a simple two arm (placebo vs. vaccine) trial with 90% power to detect a dichotomous outcome (lesion vs. no lesion) at a p value of 0.05 and with vaccine efficacy of 60%, a CHIM with 95% take rate would require 24 participants. In comparison, if the take rate was only 60%, the same study would require 74 participants. The GMP clinical parasite bank we have generated is supported by a comprehensive data package (as described here) and should be sufficient to conduct CHIM studies in >1200 individuals by sand fly transmission. Whilst manufacture was not of the scale associated with development of for example virally-vectored vaccines, it balances yield with the desire to limit in vitro parasite expansion and should serve as a key resource for several years to come.

This study has some limitations. *L major* was chosen as the challenge agent as this represents the species with most limited clinical severity. Whilst the use of a *L major* CHIM would clearly inform the development of prophylactic vaccines against Old World CL, the degree to which this data could be extrapolated to protection against other species, for example *L. donovani* a causative agent of VL is untested currently. Epidemiological data has suggested ^63^ and experimental evidence supports ^64–66^ some degree of cross-protection between parasites causing CL and VL, including following vaccination. It is reasonable to suggest therefore that successful protection following vaccination in a *L. major* CHIM would provide highly encouraging albeit not definitive evidence to support the development of vaccines against VL or other forms of CL.

Further steps in the development of this CHIM will be reported elsewhere, including the results of a study to optimise the human biting protocol using uninfected sand flies (www.clinicaltrials.gov; NCT03999970) and the results of our patient and public involvement activities. Our next step is to seek ethical approval to conduct pilot studies of human challenge using the parasite isolate described here, in order to ascertain the frequency of take and rate of lesion development in human volunteers following infectious sand fly bite, leading to appropriately designed clinical trials employing human challenge as a measure of vaccine efficacy. These pilot studies will also provide for detailed mechanistic insights into the early evolution of a primary CL lesion.

## Methods

### Ethics statement

All human studies were conducted in accord with the Declaration of Helsinki. Ethical approval was obtained from the Helsinki Committees of Hebrew University (0400-18-SOR) and The Chaim Sheba Medical Centre (5658-18-SMC) and the University of York Dept. of Biology Ethics Committee. Informed consent was obtained from patients with PCR-confirmed leishmaniasis for parasite isolation and subsequent use of these parasites in the development of a human challenge model. Animals were maintained and handled at Charles University and the University of York in accordance with institutional guidelines and national legislation (Czech republic: Act No. 246/1992 and 359/2012 coll. on Protection of Animals against Cruelty in present statutes at large; UK: Animals (Scientific Procedures) Act 1986). All the experiments were approved by: i) the Committee on the Ethics of Laboratory Experiments of the Charles University in Prague and were performed under permit from the Ministry of Education, Youth and Sports of the Czech Republic (MSMT-28321/2018-6), and ii) The University of York Animal Welfare and Ethics Review Board and performed under Home Office license (PPL P49487014).

### Mice and parasites

Adult specific pathogen free BALB/c mice were used in all experiments reported here and were obtained from either AnLab s.r.o (Prague) or Charles River UK (York). Mice were maintained in individually ventilated cages with food and water ad libitum and a 12 h light/12 h dark photoperiod. Two new clinical isolates, *L. major*_MRC-01 and *L. major*_MRC-02, were isolated from patient lesion smears by culturing in 1ml Schneider’s *Drosophila* medium (Cat. No. 21720024, Gibco) containing 20% fetal bovine serum (Australian origin, Cat. No. 10101145, Lot No. 1951998S, Gibco) at 26°C. Antibiotics were not included in the medium. The cultures were positive for promastigotes after 9 and 13 days, respectively. The parasites were further expanded and cryopreserved after one and two passages. For freezing, parasites (2 × 10^7^ cells/vial) were suspended in Schneider’s *Drosophila* medium containing 30% fetal bovine serum and 6.5% DMSO and transferred to a Mr Frosty box at −80°C.

### Parasite sequencing and analysis

DNA from promastigote cultures was extracted using a DNeasy column according to manufacturer’s instruction (Qiagen). PCR-free library preparation (Lucigen) and NextSeq 500 sequencing (Illumina) was performed at Genome Quebec. Raw reads were processed as previously described ^67,68^. Briefly, the reads were aligned to the *L. major* Friedlin reference genome sequence obtained from TryTripDB ^69^ using the Burrows Wheeler Aligner ^70^ and analysed using VarScan2 ^71^. For phylogeny generation, additional sequences obtained from GenBank from whole genome sequencing projects of *L. major* were also processed and aligned along with MRC-01 and MRC-02 isolates. Polymorphisms and copy number variant were plotted using circos ^72^ and inspected manually using the Integrative Genomics Viewer^73^.

*L. major* isolates were tested for the presence of LRV2 by RT-PCR using the primers LRV F-HR (5’-tgt aac cca cat aaa cag tgt gc-3’) and LRV R-HR (5’-att tca tcc agc ttg act ggg-3’) essentially as described by ^74^. RNA was purified from *L. major*_MRC-01, *L. major*_MRC-02 or *L. aethiopica* (MHOM/ET/1985/LRC-L494) using the TRI reagent (Sigma-Aldrich) according to the manufacturer’s instructions. The latter isolate was used as a positive control for LRV02. cDNA was synthesis using the Transcriptor Universal cDNA Master Kit (Sigma-Aldrich) with random hexamer primers. Each PCR reaction (25μl) contained 5μl cDNA, 10μM each primer and 10μl master mix (PCR-Ready High Specificity, Syntezza Bioscience); and were carried out as follows: Initial denaturation 95°C for 2 min, 35 cycles at 95°C for 20 s, annealing at 55°C for 40 s, extension at 72°C for 40 s and final extension at 72°C for 5 min. Amplicons were analysed on 1.5% agarose gels.

### *In vivo* infectivity by needle challenge

BALB/c mice were infected s.c. in the shaved rump with 100ul saline containing 10^6^ metacyclic promastigotes, selected from stationary phase cultures using PNA agglutination ^46^. Lesion development was monitored every two-three days until patency and daily thereafter. Measurements were performed in two directions using a dial caliper and the mean (8mm) and maximum single (10mm) diameter used to evaluate when mice had reached their clinical end point. For drug treatments, mice reaching a pre-determined cut-off of 4mm were randomized (n=9-10 per group) to receive either saline or paromomycin (50mg/kg, i.p daily for 10 days). Treated mice were killed at day 10 post treatment for evaluation of parasite load by qPCR for kinetoplastid DNA (see below).

### Sand fly colonies and sand fly infections

The colonies of *P. duboscqi* and *P. papatasi* (originating in Senegal and Turkey, respectively) were maintained in the insectary of the Department of Parasitology, Charles University in Prague, under standard conditions (26°C on 50 % sucrose, humidity in the insectary 60-70% and 14 h light/10 h dark photoperiod) as described previously ^75^. The sand fly colonies have been screened by RT-PCR and found to be negative for Phleboviruses (including Sandfly Fever Sicilian Virus group, Massilia virus and Toscana Virus) and Flaviviruses (targeting a conserved region of the NS5 gene).

Promastigotes from log-phase cultures (day 3-4 in culture) were washed twice in saline and resuspended in heat-inactivated rabbit blood at a concentration of 1 × 10^6^ promastigotes/ml. Sand fly females (5-9 days old) were infected by feeding through a chick-skin membrane (BIOPHARM) on the promastigote-containing suspension. Engorged sand flies were separated and maintained under the same conditions as the colony. On day 3, 6 and 15 post bloodmeal (PBM) sample sand flies were dissected and digestive tracts examined by light microscopy. Five locations for *Leishmania* infection were distinguished: endoperitrophic space (ES), abdominal midgut (AMG), thoracic midgut (TMG), cardia (CA) and the stomodeal valve (SV). Parasite loads were estimated by two methods: i) infections were qualitatively assessed in situ as light (< 100 parasites per gut), moderate (100–1000 parasites per gut) and heavy (> 1000 parasites per gut) ^76^; ii) infections were quantitatively assessed by transferring each gut into 100 μl of 0.01% formaldehyde solution, followed by homogenization and counting using a Burker chamber. *Leishmania* with flagellar length < 2 times body length were scored as procyclic forms and those with flagellar length >2 times body length as metacyclic forms ^77^.

### Sand fly to mouse transmission experiments

For transmission experiments, BALB/c mice were anaesthetized with ketamin and xylazine (62 mg and 25 mg/kg). Sand flies infected for 15 days (as above) were placed into small plastic tubes covered with the fine mesh (10 females per tube) and the tubes were held on the ear pinnae of anaesthetized mice for one hour. Engorged sand fly females were immediately dissected for microscopical determination of infection status (as described above). One group of mice was euthanized immediately post transmission and a second group of mice was followed for a period of 6 weeks p.i.

### Determination of parasite load in tissues

Sand fly-exposed ear pinnae were dissected and stored at −20°C. Extraction of total DNA was performed using a DNA tissue isolation kit (Roche Diagnostics, Indianapolis, IN) according to the manufacturer’s instructions. Lesions from needle challenge were dissected and stored at −80°C. Extraction of total DNA was performed using DNeasy tissue isolation kit (Qiagen) according to the manufacturer’s instruction. Parasite quantification by quantitative PCR (qPCR) was performed in a Bio-Rad iCycler & iQ Real-Time PCR Systems using the SYBR Green detection method (SsoAdvanced™ Universal SYBR® Green Supermix, Bio-Rad, Hercules, CA). Primers targeting 116 bp long kinetoplast minicircle DNA sequence (forward primer (13A) 5’- GTGGGGGAGGGGCGTTCT -3’ and reverse primer (13B) - 5’- ATTTTACACCAACCCCCAGTT -3’) were used ^78^. One microliter of DNA was used per individual reaction. PCR amplifications were performed in triplicates using the following conditions: 3 min at 98°C followed by 40 repetitive cycles: 10 s at 98 °C and 25 s at 61 °C. PCR water was used as a negative control. A series of 10-fold dilutions of *L. major* promastigote DNA, ranging from 5×10^3^ to 5×10^−2^ parasites per PCR reaction was used to prepare a standard curve. Quantitative results were expressed by interpolation with a standard curve. To monitor non-specific products or primer dimers, a melting analysis was performed from 70 to 95 °C at the end of each run, with a slope of 0.5 °C/c, and 5 s at each temperature.

### Statistical analysis

Data are plotted using violin plots and mean, median, 95% CI and ranges are shown as appropriate. All statistical analysis was performed with the statistical software package SPSS version 23 or with GraphPad Prism 8 for macOS (v8.4.2). Normality was evaluated using the D’Agostino-Pearson test and differences in parasite numbers in mice and sand fly tissues were tested by non-parametric (Mann-Whitney U, Mood’s median test) or parametric tests (student’s t test or ANOVA) depending on data distribution. Time to event analysis was conducted using the log-rank (Mantel-Cox) test.

## Acknowledgments

The authors thank Shaden Kamhawi (NIAID, NIH), Hira Nagasi (FDA, NIAID), Steve Reed (ONC Bio) and Jesus Valenzuela (NIAID, NIH) for helpful discussions. This work was funded by a Developmental Pathways Funding Scheme award (MR/R014973 to PMK, CL, AL, PV and CLJ). This award is jointly funded by the UK Medical Research Council (MRC) and the UK Department for International Development (DFID) under the MRC/DFID Concordat agreement and is also part of the EDCTP2 programme supported by the European Union. PV and JS were partially supported by European Regional Development Funds (project CePaViP 16_019/0000759).

## Author contributions

HA performed *in vitro* and *in vivo* needle challenge experiments, analyzed data and generated figures, JV, BV and TB conducted parasite development studies in sand fly and sand fly transmission experiments, PL and GM conducted parasite genome analysis, ES was the clinician responsible for patient recruitment and care and contributed to the clinical protocol, EG was the project manager and data coordinator, KvB conducted *in vitro* experiments, KL developed methodology and produced the research and GMP parasite banks, CL was the sponsor’s clinical representative, wrote the clinical protocol and obtained the funding, CLJ conducted experiments, PV coordinated the sand fly experiments and PK analyzed data and produced the first draft of the manuscript. VP contributed to project design. AL, CL, CLJ, PV and PK conceived the project and obtaining funding. All authors contributed to reviewing the manuscript.

## Data and materials availability

Sequence data for *L. major*_MRC-01 and *L. major*_MRC-02 are available from GenBank (BioProject ID: PRJNA633113). Parasites produced under GMP will be available for clinical assessment of candidate *Leishmania* vaccines under an appropriate MTA.

## Supplementary Materials

*Fig. S1.* Polymorphisms between *L. major*_MRC-02 before and after mouse passage

*Fig. S2.* Gene copy number variations over chromosomes 1 to 36 (LmjF.01 – LmjF.36).

*Fig.* S3. Analysis of LRV2 presence in *Leishmania* isolates by RT-PCR.

*Fig. S4.* Rate of lesion development for *L. major*_MRC-01 and *L. major*_MRC-02 in BALB/c mice after needle challenge

*Fig. S5* Sand fly development of parasites recovered from Research Banks

*Fig. S6.* Development of lesions in BALB/c mice after exposure to infected sand fly bites.

*Table S1.* Parasite loads in *P. duboscqi* and *P. papatasi*

*Table S2.* Research Bank infections post bite in BALB/c mice

