## Supplemental Figure S1-S6 for "Characterization of a new *Leishmania major* isolate for use in a controlled human infection model"

**A**

| Chromosome | Location | Reference nucleotide | Alternative Nucleotide | Freq | Gene | Notes |
|---|---|---|---|---|---|---|
| **Post-Infection – Polymorphism Gained** | | | | | | |
| LmjF.29 | 1072606 | A | AG | 29.41% | Intergenic region | Homopolymer stretch |
| LmjF.36 | 2505068 | C | CT | 35.37% | Intergenic region | Homopolymer stretch |
| LmjF.30 | 114857 | A | ATG | 35.93% | Intergenic region | Homopolymer stretch |

**B**

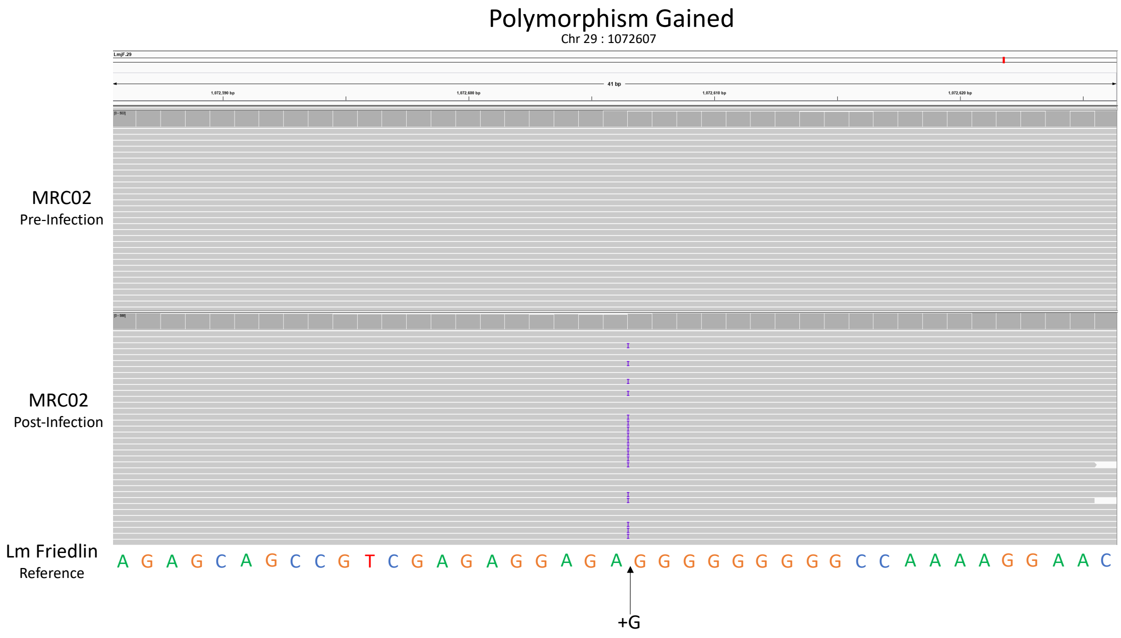

**Figure S1 Polymorphisms between *L. major*_MRC-02 before and after mouse passage**

**A.** Specific location and details of each polymorphism. Note that all the changes were in homopolymer stretches in non-coding regions and were in less that 50% of the reads.
**B.** Example of polymorphism gained on the Chromosome 29, location 1072607 where a G nucleotide was inserted in the post-infection genome in 29% of the reads as indicated.

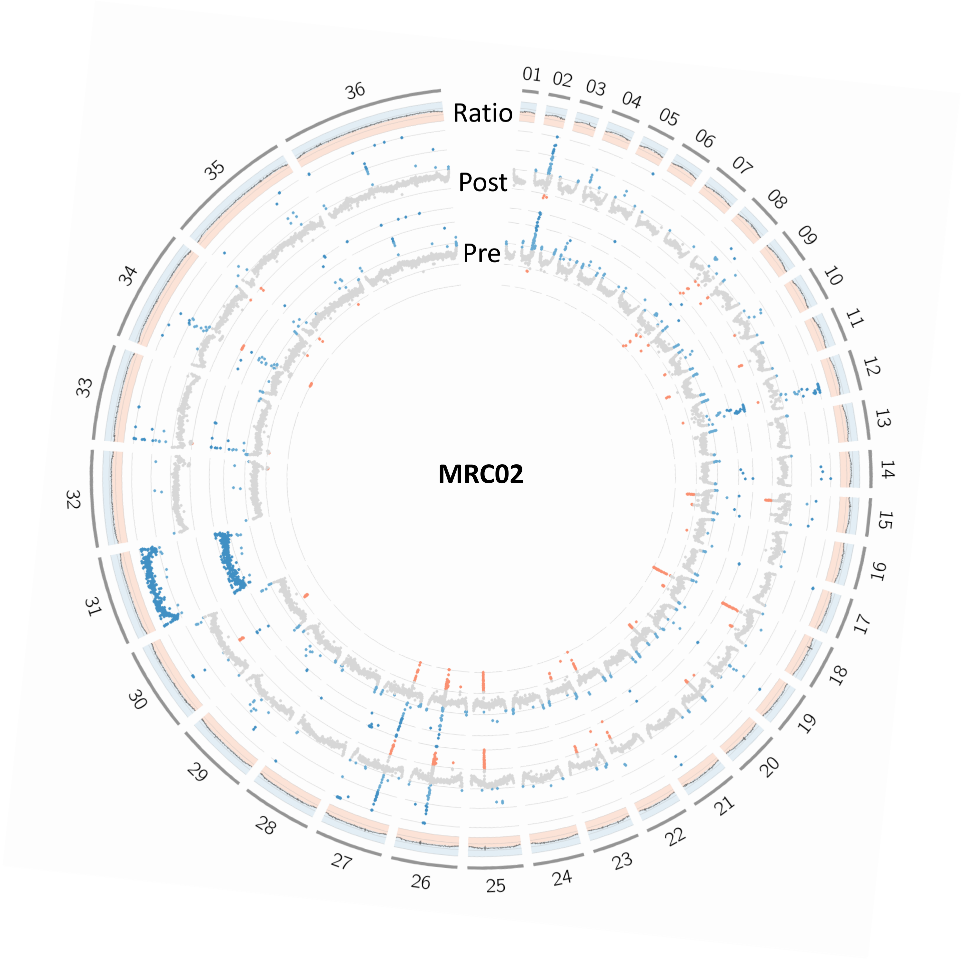

**Figure S2. Gene copy number variations over chromosomes 1 to 36.**

Raw sequencing depth per gene across all chromosomes plotted on a scale of 0-1500 and color-coded according to lower (red), median (grey), or higher (blue) read depth (Pre and Post rings). Note that chromosome 31 is tetraploid (Blue) in *L. major* while all other chromosomes are normally diploid. Coverage change ratio between the BALB/c passage (Post) and cultured (Pre) *L. major*_MRC-02 parasites shown in the outer ring on a scale of 0 to 2.

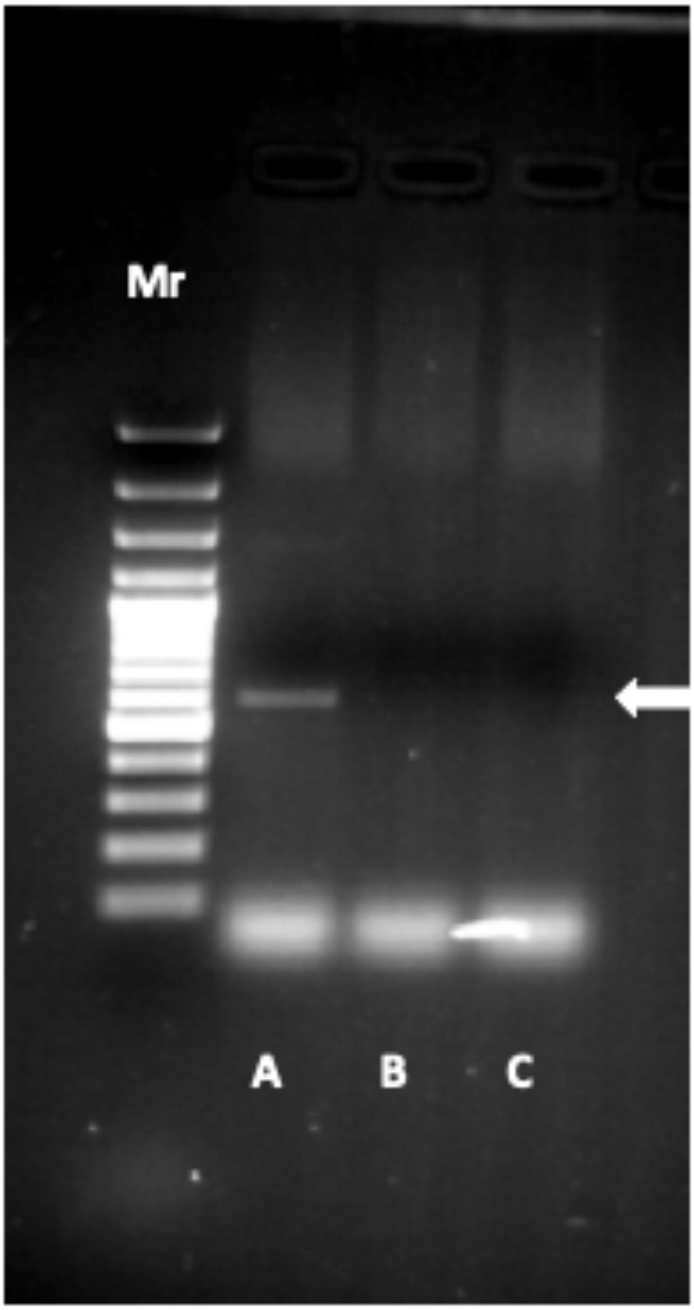

**Figure S3. Analysis of LRV2 presence in *Leishmania* isolates by RT-PCR.**

RNA was purified from each isolate and reverse transcribed into cDNA. Specific PCR targeting the RNA - dependent RNA polymerase (*RdRp*) gene of LRV2 was carried out. Arrow indicates 526 bp product. Mr - 100 bp molecular weight marker. Lane A - *L. aethiopica* LRC-L494 positive control; Lane B - *L. major*_MRC01; *L. major*_MRC02.

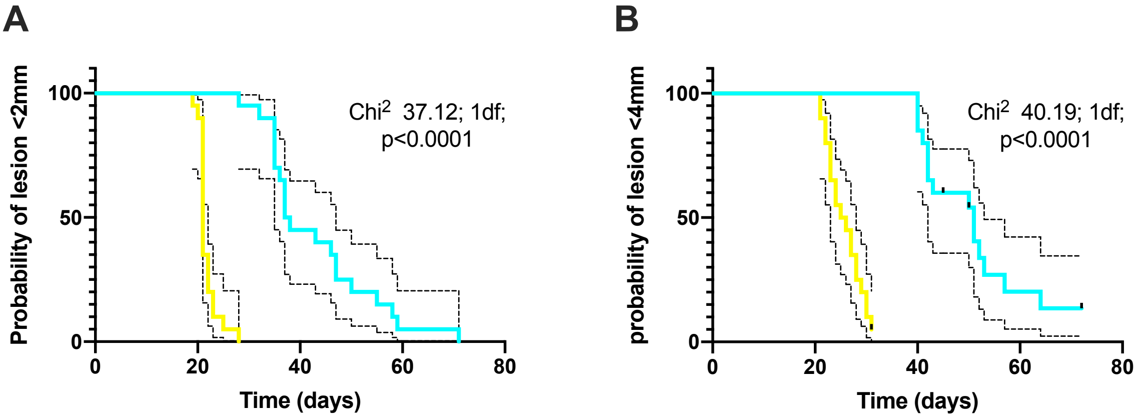

**Figure S4. Rate of lesion development for *L. major*_MRC-01 and *L. major*_MRC-02 in BALB/c mice after needle challenge**

Time to event (lesion size) is shown as Kaplan Meier plots for mice infected with *L. major*_MRC-01 (n=20, blue) and *L. major*_MRC-02 (n=20, yellow). **A**. lesion of >2mm. B. lesion of >4mm. Data were analysed using the Log-rank (Mantel-Cox) test. Dotted lines represent 95% CI. Median times to lesion > 2mm were 37.5 and 21.0 days, for *L. major*_MRC-01 and *L. major*_MRC-02 respectively (ratio 1.786, 95% CI of ratio 0.96 to 3.32; p<0.0001). Median times to lesion >4mm were 51.0 vs 25.5 for *L. major*_MRC-01 and *L. major*_MRC-02 respectively (ratio 2.0, 95% CI of ratio 1.02 to 3.94; p<0.0001).

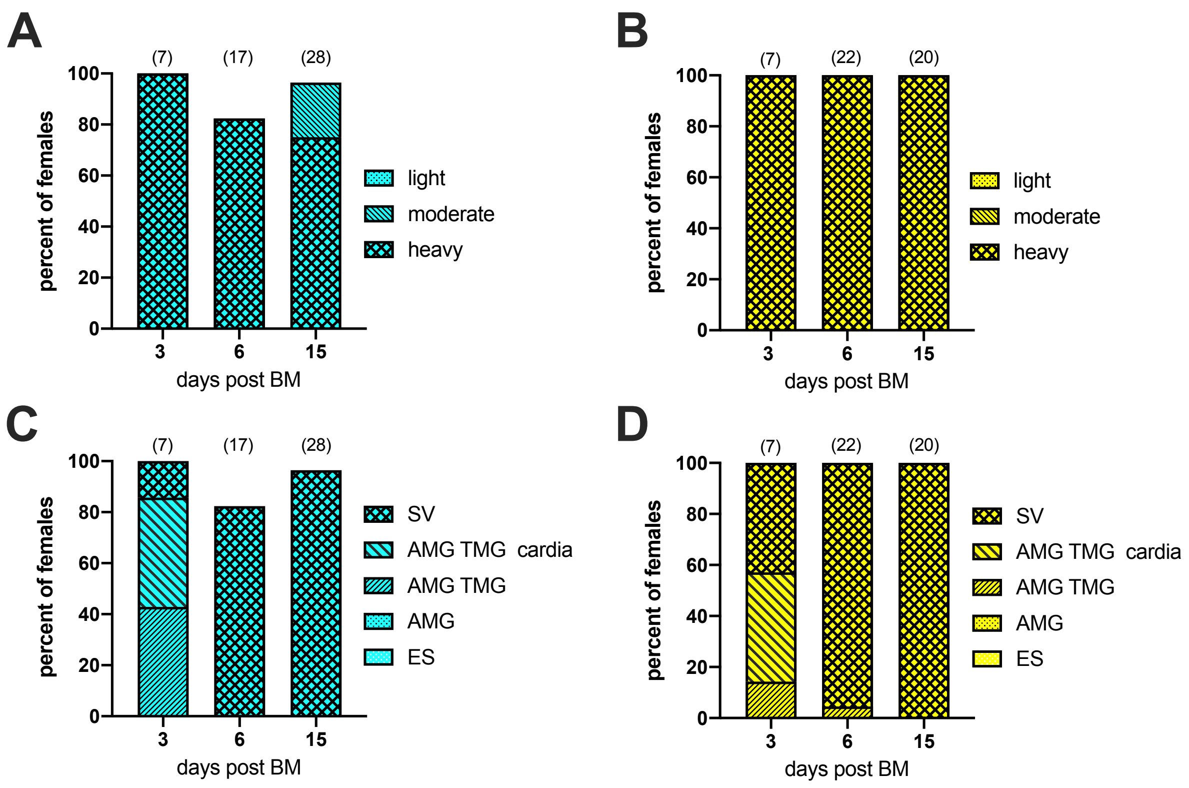

**Figure S5. Sand fly development of parasites recovered from Research Banks**

**A.** Infection rates and intensity of infections in *P. duboscqi* females infected with *L. major*_MRC-01 (1) and *L. major*_MRC-02 (2). **B.** Localisation of infections in *P. duboscqi* females infected with *L. major*_MRC-01 (1) and *L. major* _MRC-02 (2). Number of sand flies dissected is shown above each bar.

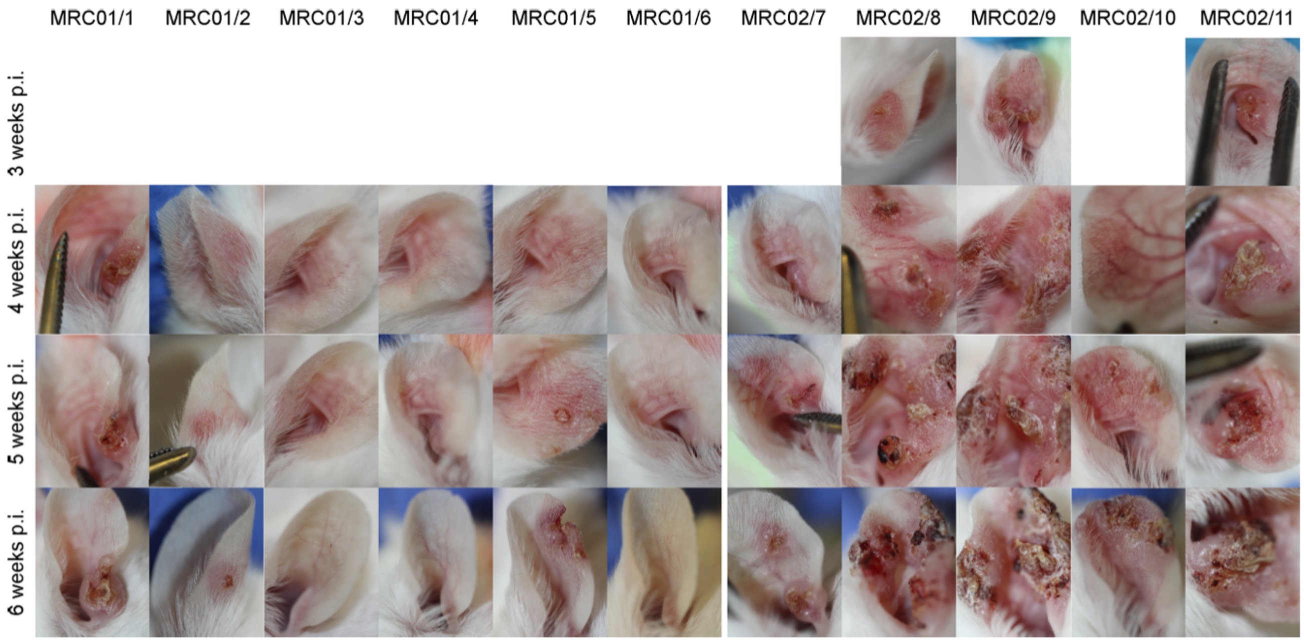

**Figure S6. Development of lesions in BALB/c mice after exposure to infected sand fly bites.**

Images show ears photographed at weeks 3-6 post exposure to 10 sand flies infected with *L. major*_MRC-01 and *L. major*_MRC-02.

### Supplementary Excel file

### Table S1. Parasite loads in *P. duboscqi* and *P. papatasi*

**See Excel table**

Sheets contain individual counts data for separated as procyclics, metacyclics or all forms for sand flies dissected at d3, d6 and d15 after an initial membrane blood meal (0 BM). Data are also shown for sand flies fed an additional blood meal at d6 (+1 BM). Summary statistics for the number of metacyclics found at day 15 PBM are also provided.

| Isolate | Sample | No sand flies | feeding rate* | No. parasite posiitve females** | No. parasites per ear | Mean no. parasites per engorged sand fly | Mean No. parasites / all infected sand flies |
|---|---|---|---|---|---|---|---|
| MRC-01 | 1 | 10 | 70% | 4+1 | 0 | 0 | 0 |
| | 2 | 10 | 60% | 2+3 | 143 | 72 | 29 |
| | 3 | 10 | 0 | 0+4 | 70 | 0 | 18 |
| | 4 | 10 | 40% | 3+2 | 7240 | 2413 | 1448 |
| | 5 | 10 | 50% | 2+4 | 506 | 253 | 84 |
| | 6 | 10 | 20% | 2+6 | 708 | 354 | 89 |
| | **Total** | **60** | **40%** | **13+20** | **8667** | **667** | **270** |
| MRC-02 | 1 | 10 | 50% | 4+2 | 3060 | 765 | 510 |
| | 2 | 10 | 40% | 1+3 | 576 | 576 | 144 |
| | 3 | 10 | 90% | 8+1 | 5670 | 709 | 630 |
| | 4 | 10 | 50% | 2+1 | 333 | 167 | 111 |
| | 5 | 10 | 70% | 1+2 | 92 | 92 | 31 |
| | 6 | 10 | 80% | 2+1 | 106 | 53 | 35 |
| | **Total** | **60** | **63%** | **18+10** | **9837** | **546** | **351** |

* Feeding rate = No. of engorged females / total No.

| *L. major isolate* | Mouse no. | No. of used sandf fly females | Feeding rate (No. of engorged females/total | No. of Leishmania positive females (fed+unfed) | First swelling appearance (weeks p.i.) | First lesion appearance (weeks p.i.) |
|---|---|---|---|---|---|---|
| MRC-01 | 1 | 10 | 60% | 4 + 2 | 3 | 4 |
| | 2 | 10 | 40% | 2 + 5 | 4 | 5 |
| | 3 | 10 | 70% | 4 + 1 | - | - |
| | 4 | 10 | 80% | 4 +0 | - | - |
| | 5 | 10 | 100% | 5 + 0 | 3 | 4 |
| | 6 | 10 | 50% | 2 + 2 | - | - |
| MRC-02 | 7 | 10 | 70% | 3 + 1 | 3 | 4 |
| | 8 | 10 | 50% | 5 + 5 | 2 | 3 |
| | 9 | 10 | 70% | 6 + 1 | 2 | 3 |
| | 10 | 10 | 70% | 5 + 2 | 3 | 3 |
| | 11 | 10 | 70% | 2 + 1 | - | 4 |

### Table S2. Research Bank infections post bite in BALB/c mice

Table shows parasite load per ear determined by qPCR post bite with 10 infected sand flies (top; sheet 1) and fly feeding rates and rate of lesion development (bottom; sheet 2).
